# Differential effects of multiplex and uniplex affiliative relationships on biomarkers of inflammation

**DOI:** 10.1101/2022.11.01.514247

**Authors:** Jessica Vandeleest, Lauren J. Wooddell, Amy C. Nathman, Brianne A. Beisner, Brenda McCowan

**Author notes:** Corresponding author details: Jessica Vandeleest, PhD.

## Abstract

Social relationships profoundly impact health in social species. Much of what we know regarding the impact of affiliative social relationships on health in nonhuman primates (NHPs) has focused on the structure of connections or the quality of relationships. These relationships are often quantified by comparing different types of affiliative behaviors (e.g., contact sitting, grooming, alliances, proximity) or pooling affiliative behaviors into an overall measure of affiliation. The influence of the breadth of affiliative behaviors (e.g., how many different types or which ones) a dyad engages in on health and fitness outcomes remains unknown. Here we employed a social network approach to explicitly explore whether the integration of different affiliative behaviors within a relationship can point to the potential function of those relationships and their impact on health-related biomarkers (i.e., pro-inflammatory cytokines) in a commonly studied non-human primate model system, the rhesus macaque (*Macaca mulatta*). Being well connected in multiplex grooming networks (networks where individuals both contact sat and groomed), which were more modular and kin biased, was associated with lower inflammation (IL-6, TNF-alpha). In contrast, being well connected in uniplex grooming networks (dyad engaged only in grooming and not in contact sitting), which were more strongly linked with social status, was associated with greater inflammation. Results suggest that multiplex relationships may function as supportive relationships that promote health. In contrast, the function of uniplex grooming relationships may be more transactional and may incur physiological costs. This complexity is important to consider for understanding the mechanisms underlying the association of social relationships on human and animal health.

## Introduction

For decades, research has shown that social relationships impact individual health and fitness in a variety of animal species, with estimates of the magnitude of the association with mortality in humans on par with other well recognized mortality risks (e.g., smoking, alcohol consumption) ^1^. While the importance of social factors for health and fitness are widely recognized, the mechanisms by which social life exerts its influence are still not well understood ^2,3^. One reason is that social life is complex; it encompasses both agonistic and affiliative interactions that vary across a range of dimensions such as frequency of interaction, symmetry, tenor, predictability, and stability ^4–6^. As such, new approaches addressing this complexity are needed to better understand the biological mechanisms by which social relationships influence health.

Traditionally, two approaches taken to unravel this complexity in humans are to examine the structural and/or functional aspects of social relationships. Structural studies frequently concentrate effort on quantifying relationships by calculating the number of social partners, the frequency of interactions, or the higher order structuring of social relationships using social network analysis ^1,3,7–13^. Functional studies attempt to examine what purpose or role a relationship serves and whether it meets the needs of the individual. In humans, functional measures of social relationships include surveys of perceived social support, informational support, emotional support, and tangible support ^1^. Both have been associated with health outcomes where some studies have found that the quantity of relationships (structural) have health benefits and others have emphasized that the types or quality of relationships (functional) are what matters (see Holt-Lunstad et al., 2010 for a meta-analysis). In comparison, for nonhuman animals, assessing the structural aspects of social relationships is common ^3,14–16^, but examining the functions of different relationships is far more challenging due to the fact we cannot directly ask animals about the value or perceptions of their relationships. Instead, research on social relationships in animals has more often relied on metrics designed to indirectly assess the quality rather than directly query the function of their relationships ^6^.

As such, studies of animal affiliation have not explicitly attempted to differentiate between structural and functional aspects of social relationships. Instead they tend to characterize the quantity and quality of relationships by measuring various affiliative behaviors, and, in doing so, these behaviors are either analyzed separately (e.g., grooming *or* proximity) or lumped together giving them roughly equal weight ^3,6^. Social network analysis is one technique well suited to studying the structure of social relationships. Typically, researchers examine the effects of individual network centrality metrics (e.g., eigenvector, betweenness, and closeness centrality) and their impact on a variety of health-related metrics. While the general pattern seen across these measures is that greater connectedness or centrality in specific behavioral networks is associated with lower risk for gastrointestinal pathogens, increased reproduction and longevity (Balasubramaniam et al., 2016; Brent et al., 2013; Cheney et al., 2016; Ostner & Schülke, 2018), there is little consistency across studies in identifying which specific social role or network metric is important, and some studies find no impact of social network position on fitness at all (Ellis et al., 2019; Ostner & Schülke, 2018; Snyder-Mackler et al., 2020). Notably, many of these networks metrics are highly correlated and, in practice, measure similar roles making the study of the underlying mechanisms challenging ^17,18^. Indeed, recent perspectives on network analysis suggest that certain metrics such as betweenness centrality only become interesting when they deviate from other network metrics such as degree centrality because if the structure of the network is a result of a few highly central individuals high betweenness is likely due to high degree ^19–21^. Thus, it is difficult to ascertain which specific metric to use and thus a suite of metrics is often employed instead ^22^.

Another commonly used metric to assess the quality of affiliative social relationships in nonhuman primates is the dyadic sociality index (DSI) which aggregates information on the quantity (frequency) of multiple, correlated affiliative behaviors (e.g., grooming and proximity). Relationships with high DSI scores are commonly referred to as strong bonds and tend to be equitable, stable, involve frequent interaction, and are most common between kin and peers ^3,23,24^. From a health perspective, higher number or quality of these strong bonds has been associated with acute hormonal responses (e.g., oxytocin or cortisol levels), increased reproduction, and longer survival ^3,9,12,25^. However, these strong affiliative bonds make up only a small fraction of the affiliative relationships an individual has ^12,23,26–28^ with weaker affiliative bonds comprising the remainder. The function of these other, weaker bonds has been hypothesized to increase social flexibility (e.g., social connections can shift with environmental demands), allowing general social integration and indirect connections that might provide access to others who may have resources or information ^3,26^. Yet, evidence for an association between weak bonds and health and fitness is mixed ^12,26^. As helpful as these metrics are, such approaches may overlook key information on the degree to which the diversity or breadth of affiliative interactions in which a dyad engages are *integrated* ^4,6^. For example, a relationship in which a specific dyad engages in multiple types of affiliation (e.g., contact sitting, grooming, *and* proximity) may differ from one in which a dyad engages in only one single behavioral domain (e.g., contact sitting only, grooming only, *or* proximity only), even if the rate of interaction is the same.

Recent advances in social network analysis and theory, and specifically multilayer or multiplex networks, (^29^; e.g., multiple types of interactions among the same set of individuals) may eventually provide a solution with tools to address such issues and, most importantly, to disentangle the impact of structural and functional social relationships. However, currently, these analytical methods (e.g., metrics like versatility and multiplex PageRank) are not sufficiently sophisticated to account for the many structural differences that exist between behavioral networks of different types (e.g., proximity networks are often dense and undirected, contact networks may be more sparse and undirected whereas grooming networks are directed)^29^. Yet, given mounting evidence for the importance of multidimensionality in social relationships, the integration of diverse affiliative behaviors at this dyadic level may provide important additional information (1) as to the nature of those relationships, (2) on how their structure points to their potential function(s), and (3) on their downstream impacts on health and fitness (Balasubramaniam et al., 2016). Indeed, in many past studies, multidimensional measures often are the strongest predictors of health and fitness ^1,8,13^. Yet, again, few animal studies to date (except see ^27^) have examined whether the integration of that diversity of interactions in a relationship provide clues as to the nature of those social relationships. One exception, conducted by Balasubramaniam and colleagues ^14^ representing one type of integrated approach, found that highly connected rhesus macaques (i.e., high outdegree or eigenvector) in a grooming network were less likely to have *Shigella*, a gastrointestinal pathogen, *but only if they were also well connected in a huddling network* (i.e., high betweenness), suggesting that the presence of multiple affiliative behaviors across an individual’s dyadic connections (e.g., integration) confer a social buffering effect on individual health (Balasubramaniam et al., 2016).

Here we employed a social network approach to explicitly explore whether the integration of diverse affiliative behaviors within a relationship can point to the potential function of those relationships and their impact on health-related outcomes in a commonly studied non-human primate model system, the rhesus macaque (*Macaca mulatta*). We use rhesus macaques as a group-living, nonhuman primate (NHP) model because their social relationships are highly differentiated, exhibit a high degree of complexity and individual variability, and have been linked to a variety of health and fitness outcomes ^30–33^. Affiliation in rhesus macaques, as in many primate species takes many forms, including grooming, contact sitting, proximity, embracing, and less commonly coalitionary support ^34–36^. In macaques, grooming is commonly used to indicate the presence of an affiliative relationship ^37,38^. Grooming has been proposed to serve multiple social functions including: to maintain social bonds ^37^ and social cohesion ^39^, and in exchange for tolerance from dominants, for agonistic support, or for access to resources ^40–42^. Although less commonly studied, contact sitting behavior (similar to huddling) may also be an important indicator of strong affiliative relationships ^43^, particularly those that may offer social buffering ^14^.

Therefore, in this study, an integration of affiliative behaviors was conducted by filtering these behavioral networks (grooming and contact sitting) into “multiplex” and “uniplex” networks. We use the term “multiplex” to refer to networks in which edges are represented by the co-occurrence of multiple affiliative behaviors and the term “uniplex” to refer to networks in which edges represent only one specific type of interaction (e.g. grooming *or* contact sitting, not both)^4,8^. Our multiplex networks were defined as the set of dyads who engaged in both grooming and contact sitting with edge-weights reflecting grooming (multiplex grooming) or contact sitting (multiplex contact sitting) frequency, while our uniplex networks were defined as the set of dyads who engaged solely in one behavior with edge weights reflecting the frequency of interaction (uniplex grooming or uniplex contact sitting). Our rationale for constructing multiplex and uniplex networks in this manner was inspired by Balasubramaniam ^14^ which identified the combination of grooming and huddling behavior to be especially protective against Shigella infection. Further, this distinction could also be hypothesized to relate to the different functions of grooming (social bonding/cohesion vs. exchange for tolerance and resources). Therefore, our primary focus for this study was to examine the differences between multiplex grooming and uniplex grooming networks. However, we also present results comparing additional complementary networks (multiplex contact sit vs. uniplex contact sit and grooming vs. contact sitting) to fully explore the impact of filtering networks in this way. Our analysis had two main goals: 1) determine if the network structure of related networks differed, and 2) the effects of individual network centrality on health. To this end, differences in global network structure of multiplex and uniplex or grooming and contact sitting networks were compared and individual-level centrality metrics from these networks were examined for their association with biomarkers of inflammation (i.e., serum pro-inflammatory cytokines), which are common, well-established indicators of individual health status ^33,44^. As such, our prediction was that individuals exhibiting more central roles in multiplex (socially cohesive) networks would show lower levels of inflammatory cytokines than those exhibiting more central roles in uniplex networks.

## Materials and Methods

### Study system

Rhesus macaques live in large multi-male, multi-female social groups organized by rank and kinship ^45^. For females, rank is inherited from their mothers and generally all members of a matriline hold adjacent ranks ^46^ (although see ^47^). In contrast, males generally immigrate into a new social group and may enter at the bottom of the hierarchy, queueing for rank, or attain rank through direct competition ^48^. Rhesus macaque females form the core of the social group with affiliation between both kin and non-kin playing a key role in maintaining group stability ^45,49^. Although male social bonds have important fitness outcomes in macaques generally ^50^, male rhesus macaques engage in social affiliation far less frequently ^51^ and tend to be more socially isolated than females ^52^. Therefore, we focused our study on females, which we predict will be more strongly impacted by social bonds than males. We use rhesus macaques as a group-living, nonhuman primate (NHP) model because their social relationships are highly differentiated, exhibit a high degree of complexity and individual variability, and have been linked to a variety of health and fitness outcomes ^30–33^.

### Subjects and housing

Subjects were 248 breeding age (3 years and older) female rhesus macaques (*Macaca mulatta*) that were born at the California National Primate Research Center in Davis, California (Table 1). Subjects lived in one of four large multigenerational and matrilineal social groups containing 100-200 mixed-sex individuals (Table 1), each housed in a 0.2 hectare outdoor enclosure. Subjects were fed commercial monkey chow and foraging enrichment twice daily. Fruits or vegetables were provided weekly. Water was available ad libitum.

**Table 1:** Group Demographics.

| **Group** | **Group size**<br>**(adults)** | **N (adult**<br>**females)** | **# of**<br>**Matrilines<sup>a</sup>** | **Mean Matriline<sup>a</sup>**<br>**size (SD)** |
| --- | --- | --- | --- | --- |
| Group A | 131 (101) | 74 | 13 | 5.7 (3.6) |
| Group B | 204 (101) | 67 | 33 | 2.0 (1.0) |
| Group C | 125 (55) | 39 | 6 | 6.5 (3.9) |
| Group D | 185 (96) | 68 | 13 | 5.2 (2.3) |

<sup>a</sup> Number of matriline and matriline size statistics include only breeding age females. Individuals were considered part of the same matriline if they could be traced back to the same female common ancestor at the time of group formation.

### Behavioral data collection

Subjects were part of a larger study on the associations between social networks and health. Groups A and B were studied for six continuous weeks during the birthing season from March to April 2013 and 2014, respectively. Groups C and D were studied for six continuous weeks during the breeding season from September to October 2013 and 2014, respectively. Behavioral data were collected six hours per day, four days per week from 0900-1200 and 1300-1600 each day by three observers (inter-rater reliability, Krippendorff’s alpha ≥0.85). Affiliative behavior was collected by one observer via scan sampling every 20-minutes (maximum 18 scans per day), where identities of all adult female dyads affiliating (i.e. grooming or contact sitting) were recorded^14^. All animals had tattoos and fur markings that allowed accurate individual identification. All observers demonstrated animal ID reliability of > 95%. Grooming was defined as cleaning or manipulating the fur of another animal and contact sitting included ventral contact, embrace, or side by side sitting for at least 3 seconds. During each scan, these behaviors were mutually exclusive for a dyad (an individual grooming another was not contact sitting with that individual). Affiliation scans produced 1637 scans (Group A: N=418, Group B: N=410, Group C: N=378, Group D: N=431) and a median of 38 grooming interactions per female (group range 23 – 49) and 28 contact sitting interactions (group range 13-52). This sampling scheme has been shown to produce sufficiently sampled grooming and contact sitting networks^38^. Aggression data (threats, chases, bites) were collected via an event sampling protocol for six hours per day, four days per week by two other observers (average of 42.5 interactions per individual, group range 36.2 – 51.9). Because social status has been shown to impact inflammation^31^ (although see^30^), dyadic aggression data was used to calculate dominance ranks and dominance certainty via the R package *Perc*^30,53^. Dominance rank was expressed as the percent of animals in the group outranked and therefore ranged from 0 (low) to 1 (high).

### Affiliative network analysis

First, weighted networks were constructed from grooming and contact sitting interactions (Figure 1A). Each of these networks (i.e., grooming or contact sitting) were then separated into two more networks, a multiplex network where edges between dyads that both groomed and contact sat were retained (Figure 1C or 1E), and a uniplex network in which edges were retained for dyads that only groomed (Figure 1D) or only contact sat (Figure 1B). Edge-weights in contact sitting networks (all contact sitting, uniplex contact sitting, multiplex contact sitting) reflected the number of unique scans in which a dyad was observed contact sitting over the 6-week period. Edge-weights in grooming networks reflected the number of unique scans a dyad was observed grooming (Table S1). For each of the 6 networks (all grooming, multiplex grooming, uniplex grooming, all contact sitting, multiplex contact sitting, uniplex contact sitting), centrality and cohesion measures for each individual were calculated in R (ver. 4.0.5) using iGraph (ver. 1.3.0). The effects of the direct connections for individuals were measured using degree centrality and strength. The effect of an individual’s indirect connections in the network was evaluated using eigenvector, betweenness, and closeness centralities ^3,14,16^. In addition, the degree to which individuals were part of cohesive local communities was measured by the local clustering coefficient (i.e., triadic closure). Multiple metrics were chosen to reflect the different ways social integration can manifest (e.g., bridging, cohesion, embeddedness, etc. Table 2).

**Figure 1:**
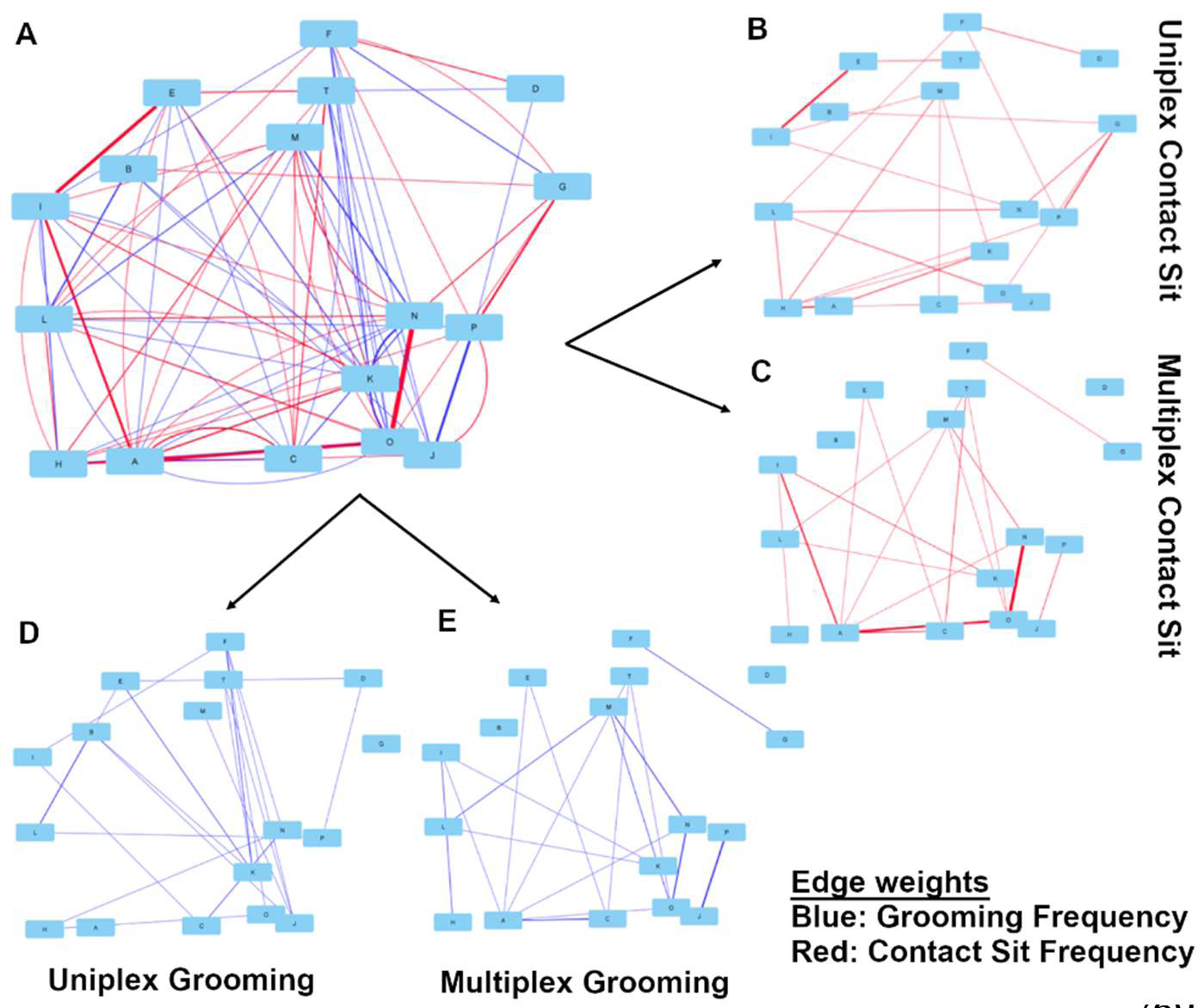
A network example using a subset of animals from Group C. Edge weight is represented using line thickness and behavior is indicated by color. Network filtering started with networks consisting of (A) all observed grooming (blue) and contact sitting (red) interactions. Edges from these networks were then filtered into multiplex grooming (E) or multiplex contact sitting (C) networks if dyads engaged in both behaviors. Dyads that only groomed were filtered into a uniplex grooming network (D) and dyads only engaged in contact sitting were filtered into a uniplex contact sitting network (B).

**Figure 2.**
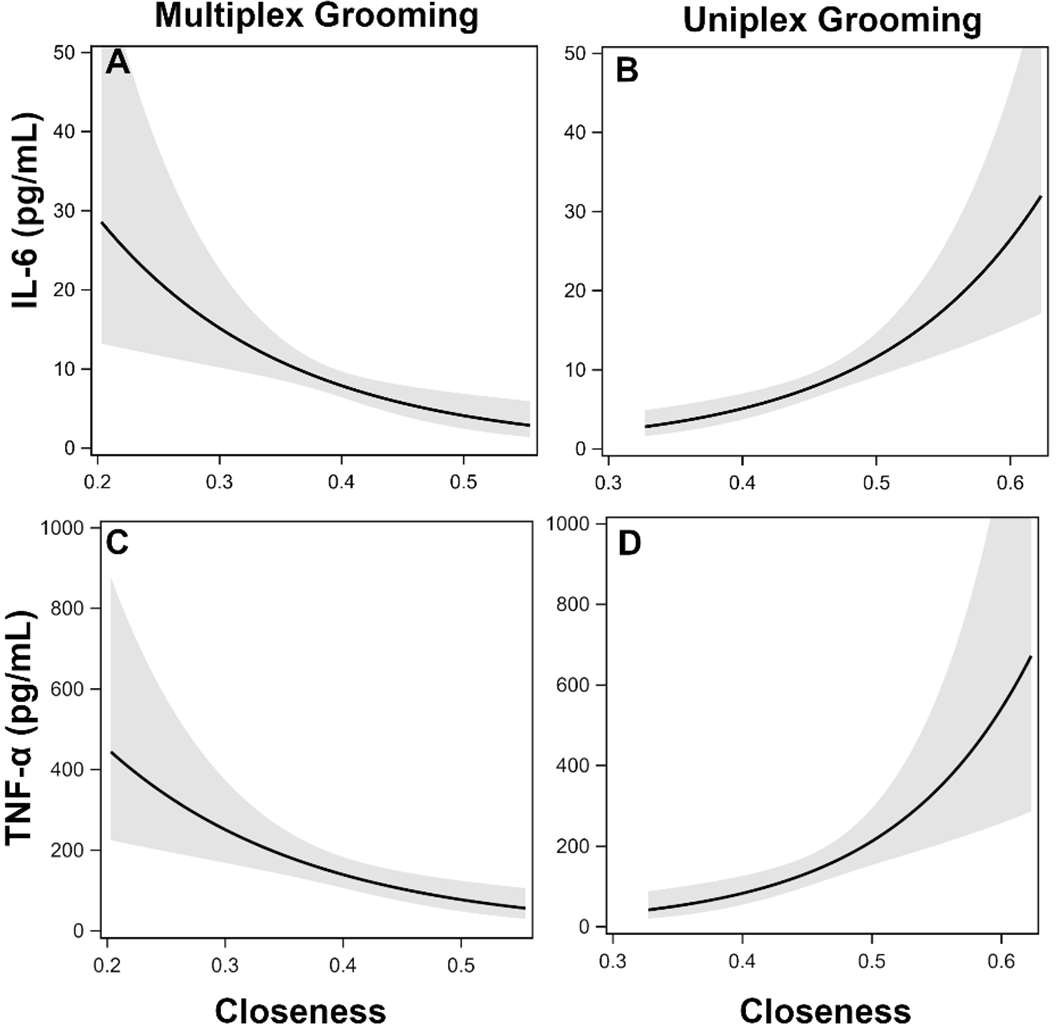
Effects of affiliative centrality on cytokines. Effects of multiplex (Multi) grooming closeness (A) and uniplex (Uni) grooming closeness (B) on levels of IL-6 with 95% confidence intervals (Model 1). Effects of multiplex grooming closeness (C) and uniplex grooming closeness (D) on levels of TNFα with 95% confidence intervals (Model 1).

**Table 2:** Network Metric Definitions.

| Measure | Description |
| --- | --- |
| Degree/Strength | measures the *number* (unweighted) of partners or *frequency* of interaction (i.e., strength) for each node.<sup>54</sup> |
| Eigenvector | measures whether individuals are well connected to others that are also well connected, a measure of *social capital*.<sup>54</sup> |
| Betweenness | measures the number of times a node lies on the shortest path between other nodes, which reflects an individual's role in connecting others in the network or acting as a *social bridge*.<sup>54</sup> |
| Closeness | measures how close each node is to all other nodes within the network, which reflects how *embedded* an individual is in the network.<sup>55</sup> |
| Clustering Coefficient | measures the extent to which a node's neighbors are also connected to each other, a measure of *cliquishness* or *subgrouping*.<sup>56</sup> |

### Biological sample collection

Blood samples were taken during the fifth week of each group’s study period during routine, semi-annual health checks. On a single morning, all animals in a group were lightly sedated with ketamine (10 mg/kg) and given veterinary exams. Blood samples were obtained from the femoral vein and serum was aliquoted and stored at −80 °C for later assay. The order in which animals were processed and samples were collected was recorded to control for any potential impacts of the sampling procedure on the physiological variables examined.

### Pro-inflammatory Cytokines

Chronic inflammation is associated with a variety of diseases (e.g., diabetes, cardiovascular disease, cancer) and mortality ^57–59^, and high levels of pro-inflammatory cytokines, such as IL-6 and TNF-α, have previously been reported to be associated with social variables (e.g., low social status, low social integration, poor quality relationships, loneliness) in both humans and rhesus macaques ^30,33,44,60,61^. Therefore, we chose to measure serum levels of IL-6 and TNF-α as a general biomarker of health. Serum levels of IL-6 and TNF-α were measured simultaneously using commercially available, species specific Milliplex multi-analyte profiling (MAP) reagents purchased from EMD/Millipore (Billerica, MA, USA), and utilizing Luminex Xmap technology (Luminex, Austin, TX, USA). Color coded polystyrene microbeads coated with specific antibodies for IL-6 and TNF-α were incubated with the serum samples, washed, and then further reacted with biotinylated detector antibodies followed by Streptavidin-PE to label the immune complexes on the beads. After a final washing to remove all unbound material, the beads were interrogated in a BioPlex dual laser (BioRad, Hercules, CA, USA). The median fluorescent index for each sample was compared to a standard curve to calculate the concentration (IL-6: mean = 12.55 pg/mL, sd = 46.92, range = 0 – 690; TNF-α: mean = 185.0 pg/mL, sd = 442.27, range = 0 – 4052; see Figure S2 for histograms). Samples were tested in duplicate and had an intra-assay coefficient of variability of 15.3%. Samples were re-analyzed if the CV was greater than 25% for all analytes measured. Manufacturer provided quality control samples fell within recommended ranges for all assays. Samples falling below the threshold sensitivity of the assay (1.6 pg/mL) were assigned a value of zero (IL-6: N = 77, TNF-α: N = 56).

### Statistical analysis

Two sets of analyses were done to determine whether 1) multiplex and uniplex grooming networks (i.e., Fig 1B vs 1C or Fig. 1D vs 1E) or grooming and contact sitting networks (Fig. 1A red vs. blue) differ in structure and relationships to known social features of rhesus macaques (e.g., kin bias, hierarchical organization), and 2) whether network metrics from these networks predicted biomarkers of inflammation, with a specific focus on the relative impact of multiplex vs uniplex (or grooming vs. contact sitting) network position on inflammation.

First, to validate this method, the structure of the multiplex and uniplex networks, which were treated as weighted and directed (grooming only) networks, were compared to determine if they exhibited differences in key structural features of rhesus relationships. For example, evidence suggests that despotic macaques such as rhesus, particularly in large groups, are likely to have grooming networks that are modular (i.e., shows subgrouping), expected to be based on kinship, and have individual network positions (i.e., eigenvector centrality) that are correlated with rank^62,63^. Therefore, we examined whether these two networks differed in the degree of clustering (Newman’s modularity, clustering coefficient), kin bias (e.g., proportion of kin (kin unweighted degree/total unweighted degree)), and associations with rank (proportion of grooming up the hierarchy, rank disparity among grooming dyads) for each of the four groups studied. Also, because previous research has focused on bond strength, we further examined reciprocity, strength of relationships (average edge weight), and distribution of grooming (eigenvector centralization) across these network types. Due to the low number of groups in the comparison, paired t-tests were used to evaluate if network metrics were consistently different across groups. Normality of the differences was evaluated using the Shapiro-Wilk test, and if significant then Wilcoxon signed rank tests were used. Grooming and contact sitting networks were also compared. As a final structural analysis, we examined the correlations between individual level network positions from these two network types (Table S2) to evaluate multicollinearity within networks and associations between networks, including contact sitting networks.

Next, to determine if an individual’s position in the multiplex or uniplex networks was associated with pro-inflammatory cytokines, we ran generalized linear models using a negative binomial distribution (R package lme4 v.1.1-34) on each biomarker separately (see ^30^ for details on distribution choice and Figure S2 for distributions). For these analyses, networks were treated as weighted but undirected because of our focus on the qualities of a relationship rather than focused specifically on grooming behavior and because contact sitting is recorded as an undirected behavior (i.e., information on who initiated the interaction is unavailable). One animal was excluded from the IL-6 analyses because it was an outlier with influence (Cook’s D >1); all other outliers had a Cook’s D < 0.5 and therefore were included in the analyses. A second animal was excluded from all analysis due to the fact she was not included in the uniplex network. Model building proceeded in five steps for each outcome (i.e., IL-6, TNF-α, see Figure S2). For all steps, ΔAIC > 2 was used to identify potential predictors and candidate models. For one social group, 6 animals were not present in the uniplex contact sit network (they never engaged in contact sitting with someone they did not also groom). Therefore to evaluate the effects of uniplex contact sitting we use a subset of the full dataset for AIC comparisons. First, a random effect indicating the group ID was evaluated for each outcome, and all subsequent models were compared to this random effects only model. Second, variables from the literature (age, dominance rank, dominance certainty, sampling order), although not of direct interest here, were evaluated to determine if it was necessary to control for their effects on inflammation before examining social network variables. Third, due to the large number of potential predictors in our exploratory analysis, a statistical winnowing strategy was used to reduce the number of social network variables under consideration ^64^. This involved running univariate models for each metric from each network (6 networks with 6 metrics each). Due to our goal of directly comparing effects of centrality in different networks and predictions for multiplex vs uniplex networks, if no metric generated improvement in model fit (i.e., ΔAIC > 2) for a given network all available network metrics for that network were explored in step 4. Fourth, to compare the effects of uniplex and multiplex network connectivity directly, candidate predictors from step 3 for the uniplex and multiplex grooming (Figure 1D vs E), uniplex and multiplex contact sitting networks (Figure 1B vs C) were directly compared. Additionally, effects of centrality in grooming vs contact sitting networks were compared (i.e., Figure 1A red vs blue). Fifth, a final set of best models was identified by comparing AIC across all models generated in steps 3 and 4. Metrics from the same network were never included in the same model due to the interdependence of network metrics. If no single best model emerged, candidate models (i.e., those with ΔAIC ≤ 2) are discussed. A log of all models tested is available in Tables S3-4.

### Ethical Note

All procedures used in this study met all legal requirements of the United States as well as guidelines set by the American Society of Primatologists regarding the ethical treatment of non-human primates. This study was approved by the Institutional Care and Use Committee at the University of California, Davis and was carried out in compliance with the ARRIVE guidelines.

## Results

### Multiplex vs. Uniplex Affiliation Networks

For all groups studied, clear differences in network topology, kinship, and associations with dominance rank were seen between the multiplex and uniplex affiliative networks (Table 3). Multiplex grooming networks had higher average edge-weight (the average number of interactions per social partner), clustering coefficient, and modularity (the degree to which the network can be divided into subgroups) than uniplex grooming networks for all groups. Notably, although average edge-weights in the multiplex networks were higher than uniplex networks, the predominant edge weight in all networks was 1-2 interactions (Figure S1). Multiplex grooming networks also consistently showed more kin bias (proportion kin) and reciprocity than uniplex grooming networks. In contrast, both multiplex and uniplex grooming networks showed associations between rank and affiliation (i.e., grooming was directed up the hierarchy and eigenvector centrality was correlated with rank in both networks) but the disparity in the ranks of the grooming partners was greater in the uniplex affiliation networks compared to the multiplex networks. Results for the multiplex vs uniplex contact sitting networks were the same as for multiplex and uniplex grooming networks with the exception of reciprocity and grooming up the hierarchy which were not calculated for these undirected networks (Table 3). Individual centrality metrics generated from the multiplex grooming networks were largely uncorrelated with metrics from the uniplex grooming networks (mean correlation strength = 0.11, SD = 0.08, Table S2). In contrast, centrality metrics from grooming and contact sitting (mean correlation strength = 0.37, SD = 0.07) and multiplex and uniplex contact sitting networks (mean correlation strength = 0.32, SD = 0.24) were moderately correlated. In contrast, the structure of the all grooming and all contact sitting networks did not differ on any examined metric with the exception of average edge-weight which was higher in grooming networks.

**Table 3:** Whole Network Metric Comparisons.

| Network | <u>Multi vs. Uni</u><br><u>Grooming</u> | <u>Multi vs. Uni</u><br><u>Contact Sit</u> | <u>Contact Sit vs</u><br><u>Grooming</u> |
| --- | --- | --- | --- |
| Density | Multi = Uni | Multi = Uni | CS = GR |
| Modularity | Multi > Uni* | Multi > Uni* | CS = GR |
| Eigenvector Centralization | Multi = Uni | Multi = Uni | CS = GR |
| Avg Edge Weight | Multi = Uni<sup>a</sup> | Multi = Uni<sup>a</sup> | CS < GR* |
| Clustering Coefficient | Multi > Uni* | Multi > Uni* | CS = GR |
| Reciprocity | Multi > Uni* | - | - |
| Proportion Kin | Multi > Uni* | Multi > Uni* | CS = GR |
| Proportion Up Rank | Multi < Uni* | - | - |
| Rank Disparity | Multi < Uni* | Multi < Uni* | CS = GR |
| Rank/Eigenvector centrality correlation | Multi = Uni | Multi = Uni | CS = GR |

Multi: Multiplex network; Uni: Uniplex network; GR: Grooming; CS: Contact Sit. Effect indicates the overall difference between networks for all groups using a paired t-test. <sup>a</sup> Wilcoxon test. \* p < 0.05

### Relationship Dimensionality and Biomarkers of Inflammation

#### IL-6

There were six models that had AIC within 2 of the best fit model, all of which included metrics from both the multiplex and uniplex grooming networks. For all models, less connected individuals in the multiplex grooming network (degree, closeness, or clustering coefficient) but more connected individuals in the uniplex grooming (strength, closeness) network had higher levels of IL-6, although these effects were not significant in all candidate models (Table 4). Due to this directional consistency we present the results of the just the best fit model in Figure 3A-B. Multiplex closeness was not correlated with uniplex closeness (r = −0.04). Contact sitting degree was weakly negatively correlated with uniplex closeness (−0.13) but more strongly correlated with multiplex closeness (r = 0.77) yet analysis of VIF (< 2.5) and tolerance (≥ 0.4) did not indicate any issues with collinearity.

**Table 4:** IL-6 Candidate Model Outputs.

| Model Parameters | Model 1 | Model 2 | Model 3 | Model 4 | Model 5 | Model 6 |
| --- | --- | --- | --- | --- | --- | --- |
| Intercept | 0.03 | 2.8** | 1.6*** | -0.7 | 1.8*** | 1.8** |
| Uni GR Strength | - | 0.05** | 0.04** | - | 0.04** | 0.04** |
| Uni GR Closeness | 7.6** | - | - | 6.2** | - | - |
| Multi GR Degree | - | - | - | - | - | -0.03 |
| Multi GR Closeness | -3.6* | -3.2 | - | - | - | - |
| Multi GR Clustering | - | - | - | - | -0.4 | - |
| $\Delta AIC$ | 0.00 | 0.30 | 0.70 | 1.40 | 1.90 | 1.90 |

\* $p < 0.05$ , \*\* $p < 0.01$ . Uni = Uniplex, Multi = Multiplex, GR = Grooming network. All models were run using a negative binomial distribution and included a random effect of group.

#### TNF-α

There were five candidate models identified; the top four of these included comparisons of multiplex and uniplex grooming centrality and the last included centrality metrics from the grooming and contact sitting networks (see Table 5). As with IL-6, lower centrality in the multiplex grooming network (degree or closeness) but higher centrality in the uniplex grooming network (strength or closeness) were consistently associated with higher levels of TNF-α (Table 5, Figure 3C-D). In the fifth candidate model, higher eigenvector centrality in the grooming network but lower closeness centrality in the contact sitting network were associated with higher levels of TNF-α.

**Table 5:**
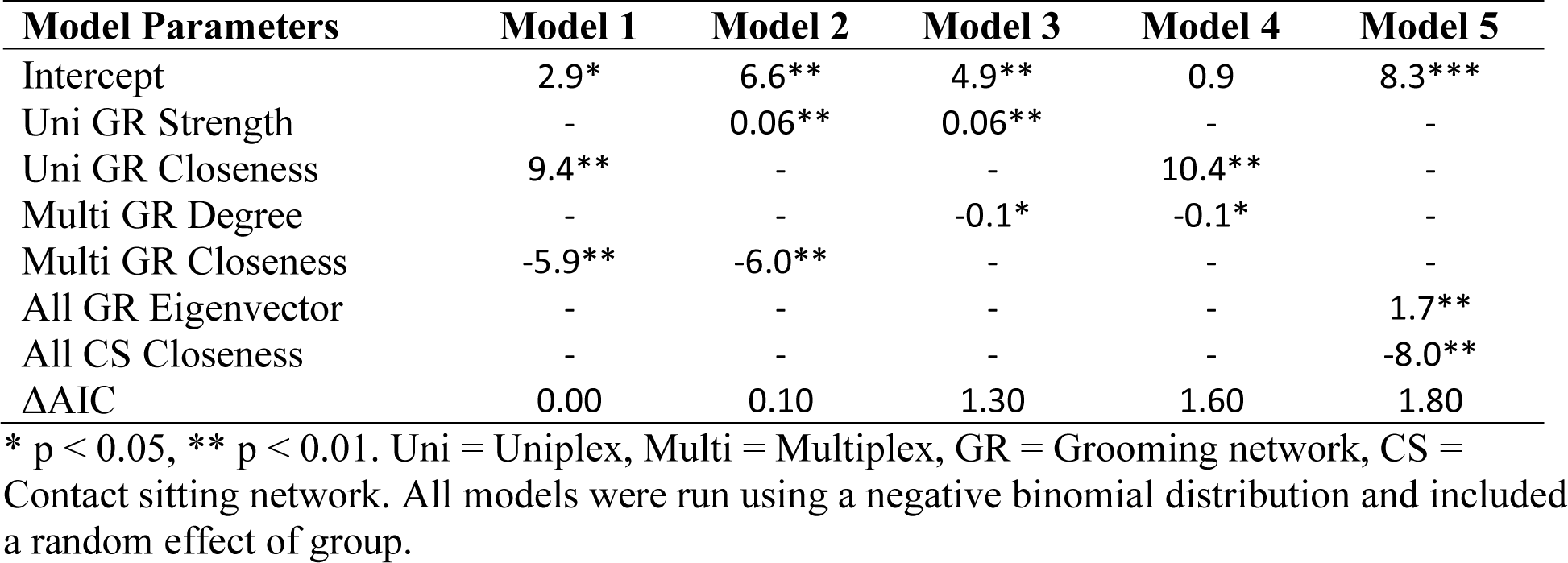
TNF-α Candidate Model Outputs.

| Model Parameters | Model 1 | Model 2 | Model 3 | Model 4 | Model 5 |
| --- | --- | --- | --- | --- | --- |
| Intercept | 2.9* | 6.6** | 4.9** | 0.9 | 8.3*** |
| Uni GR Strength | - | 0.06** | 0.06** | - | - |
| Uni GR Closeness | 9.4** | - | - | 10.4** | - |
| Multi GR Degree | - | - | -0.1* | -0.1* | - |
| Multi GR Closeness | -5.9** | -6.0** | - | - | - |
| All GR Eigenvector | - | - | - | - | 1.7** |
| All CS Closeness | - | - | - | - | -8.0** |
| ΔAIC | 0.00 | 0.10 | 1.30 | 1.60 | 1.80 |

\* $p < 0.05$ , \*\* $p < 0.01$ . Uni = Uniplex, Multi = Multiplex, GR = Grooming network, CS = Contact sitting network. All models were run using a negative binomial distribution and included a random effect of group.

## Discussion

Social primates have a complex web of differentiated social relationships, which vary in their structure and function. While strong affiliative social relationships are usually associated with better health, less is known on how the multidimensionality or integration of different affiliative behaviors within a social relationship might impact health. We identified affiliative relationships that were multiplex (animals affiliated using both grooming and contact sitting behavior) versus uniplex (animals only groomed or only contact sat). Examination of these networks revealed that they differed in topology, kinship, and associations with rank. Multiplex networks were more modular, clustered, reciprocal, had higher average edge weights, and were more strongly associated with kinship. In contrast, dyads in the uniplex networks tended to be of more disparate ranks. Notably, these differences in kinship and rank between multiplex and uniplex networks were not apparent when looking at all grooming or contact sitting interactions. The health impacts of these two networks differed as well, with females that were *less* socially embedded in multiplex grooming networks exhibiting higher levels of biomarkers of inflammation (IL-6 and TNF-α), whereas females *more* socially connected in uniplex grooming networks exhibited higher levels of biomarkers of inflammation. Notably, these effects were primarily seen in multiplex and uniplex grooming, not in the other networks tested. These results suggest that grooming which occurs in the context of multiplex affiliative relationships may result in health benefits (i.e., reduced inflammation) while grooming occurring in uniplex affiliative relationships may have potential costs.

Networks consisting of dyads with multiplex relationships showed differences from uniplex relationships in network topology, kinship, and associations with dominance. Multiplex networks had structural characteristics consistent with strong bonds or supportive affiliative relationships^23,65,66^. Specifically, interactions in the multiplex networks were more likely to be reciprocal, frequent (i.e., higher edge-weight), clustered, and associated with kinship, suggesting they are relationships that are regularly maintained and potentially more stable across time^23,66^. Previous methods demonstrating that strong bonds enhance fitness, particularly those using sociality indices, have also used multiple behaviors to assess relationship strength (e.g., grooming and proximity^12,26,27^). However, these methods rely on total duration or frequency of affiliation to describe relationships rather than characterizing the breadth or dimensionality of the relationships (e.g., dyads can have high DSI through grooming, proximity, or both). Similar to strong bonds, multiplex affiliative relationships may improve health and fitness by buffering individuals from the negative impacts of stress, improving predator detection, promoting offspring survival, and improving social stability^14,16,67,68^.

Also consistent with the literature on strong affiliative bonds, being well connected to others was associated with biomarkers of better health. Specifically, the negative association between multiplex grooming centrality (e.g., degree, closeness centrality, or clustering coefficient) and biomarkers of inflammation indicated that individuals that were generally well connected in the network (e.g., at the core of the group) may be at lower risk for inflammation related diseases^57^. Our results add to the literature suggesting that strong bonds may improve fitness by altering endocrine and immune function^13,25,69,70^. Consistent with this idea, Yang et al.^71^ found in humans that socially integrated individuals (i.e., those with more social connections across multiple domains) exhibited lower inflammation, whereas social strain (e.g., higher levels of family criticism or demands) was associated with greater inflammation. Given that familial and friend relationships tend to endure through extended periods, often persisting over decades (in both humans and NHPs), these relationships may have an important and long-lasting impact on health.

Uniplex grooming relationships may reflect relationships that are more transactional in nature ^72^. The fact that uniplex grooming relationships are less kin biased but likely to occur between dyads of more disparate ranks suggests that these relationships may be more related to grooming being used as a commodity in exchange for tolerance or support from higher ranking animals. These relationships are likely more transactional in nature, reflecting a desire to maintain peace/tolerance or used in a biological market exchange^40,41^, rather than reflecting a strong affiliative relationship. The positive association between females’ connectedness in uniplex grooming networks and biomarkers of inflammation suggests that uniplex grooming relationships may not be supportive on their own and instead are associated with increased physiological costs, at least in the short term. Specifically, predictors of inflammation in the uniplex grooming networks included strength or closeness centrality. However, the various network centrality metrics from the uniplex grooming network were more highly correlated with each other than the other networks, and therefore it is difficult to identify which specific aspect of uniplex grooming centrality might be driving these effects. However, collectively this group of candidate predictors indicates that greater general connectedness (direct and indirect) in uniplex grooming was associated with increased inflammation. Uniplex grooming relationships were maintained through generally less frequent interactions that were more likely to occur between animals of disparate ranks which may result in greater uncertainty regarding the outcome of any given interaction. This uncertainty may be stressful, and therefore have at least short-term physiological costs^73^. If these relationships are more transactional in nature, reflecting a desire to maintain peace/tolerance or used in a biological market exchange^40,41^, then maintaining more of these transactional relationships may result in increased stress, which if sustained can result in long-term physiological costs^13^. It is possible that these short-term costs are actually investments that may manifest in future benefits (e.g., tolerance, alliance support) that would offset these costs, yet this is difficult to test as the “commodities” exchanged may be heterogeneous and the time-scale for market exchanges is often unclear^74^. However, other work points to benefits of weak or economically based bonds to survival and reproduction^9,26^ (although see^12^). While these types of connections may have ultimate fitness benefits (e.g., alliance support, increased access to food), this research suggests they may also be associated with proximate costs.

## Conclusion

Both humans and many species of NHPs engage in a complex interconnected system of social interactions. Understanding the mechanisms by which social relationships impact health and fitness remains a challenge. Decades of research has established that affiliative social relationships can benefit health, however, the complexity and multidimensionality of relationships has yet to be explored. By utilizing a network approach, we were able to characterize two types of affiliative social relationships that differed in their network topology, kin bias, associations with rank, and importantly their associations with biomarkers of inflammation. Our research has indicated that features of multiplex affiliative relationships are consistent with the concept of a strong supportive relationships and may support health and fitness. In contrast, more transactional affiliative relationships (e.g., uniplex affiliation) may incur short-term health costs yet may result in ultimate benefits through commodity exchange. Still unclear is whether these effects are specific to the combination of behaviors used here (i.e., contact sitting and grooming), or if other affiliative behaviors (e.g., proximity) might provide similar information. Further research into the dimensionality of relationships might reflect different qualities or functions of relationships is needed. However, this complexity is important to consider for understanding the mechanisms underlying the impact of social relationships on human and NHP health.

## Supporting information

Dataset_NetworkLevel

Dataset_IndividualLevel

RCode_IndividualCentralityFunction

RCode_IndividualLevelAnalysis

RCode_NetworkLevelAnalysis

## Acknowledgements

We thank the data collection team: A. Barnard, T. Boussina, E. Cano, H. Caparella, C. Carminito, J. Greco, M. Jackson, A. Maness, S. Seil, N. Sharpe, A. Vitale, & S. Winkler. This research was funded by an NIH grant awarded to BM (R01-HD068335) and the California National Primate Research Center base grant (P51-OD01107-53). This is an updated version of a manuscript on the PeerJ preprint server (https://doi.org/10.7287/peerj.preprints.27961v1).

