## SupplementaryTables_Figures for "Differential effects of multiplex and uniplex affiliative relationships on biomarkers of inflammation"

### Supplementary Materials

**Jessica Vandeleest[1*], Lauren J. Wooddell[2], Amy C. Nathman[1], Brianne A. Beisner[3], Brenda McCowan[1]**

[1]California National Primate Research Center, University of California, Davis, CA, United States

[2]Department of Neurosurgery, Emory university, Atlanta, GA

[3]Emory National Primate Research Center Field Station, Lawrenceville, GA, United States

[*]Corresponding author details: Jessica Vandeleest, PhD

Contents

Figure S1: Histograms of edge weights by network and social group.

Figure S2: Cytokine histograms.

Figure S3. Flowchart of the analytical framework

Table S1: Network Node and Edge Details

Table S3: Model Building Log for IL-6

Table S4: Model Building Log for TNF-α

**Figure S1: Histograms of edge weights.** Multi: Multiplex affiliation network. Uni: Uniplex affiliation network.

Group A

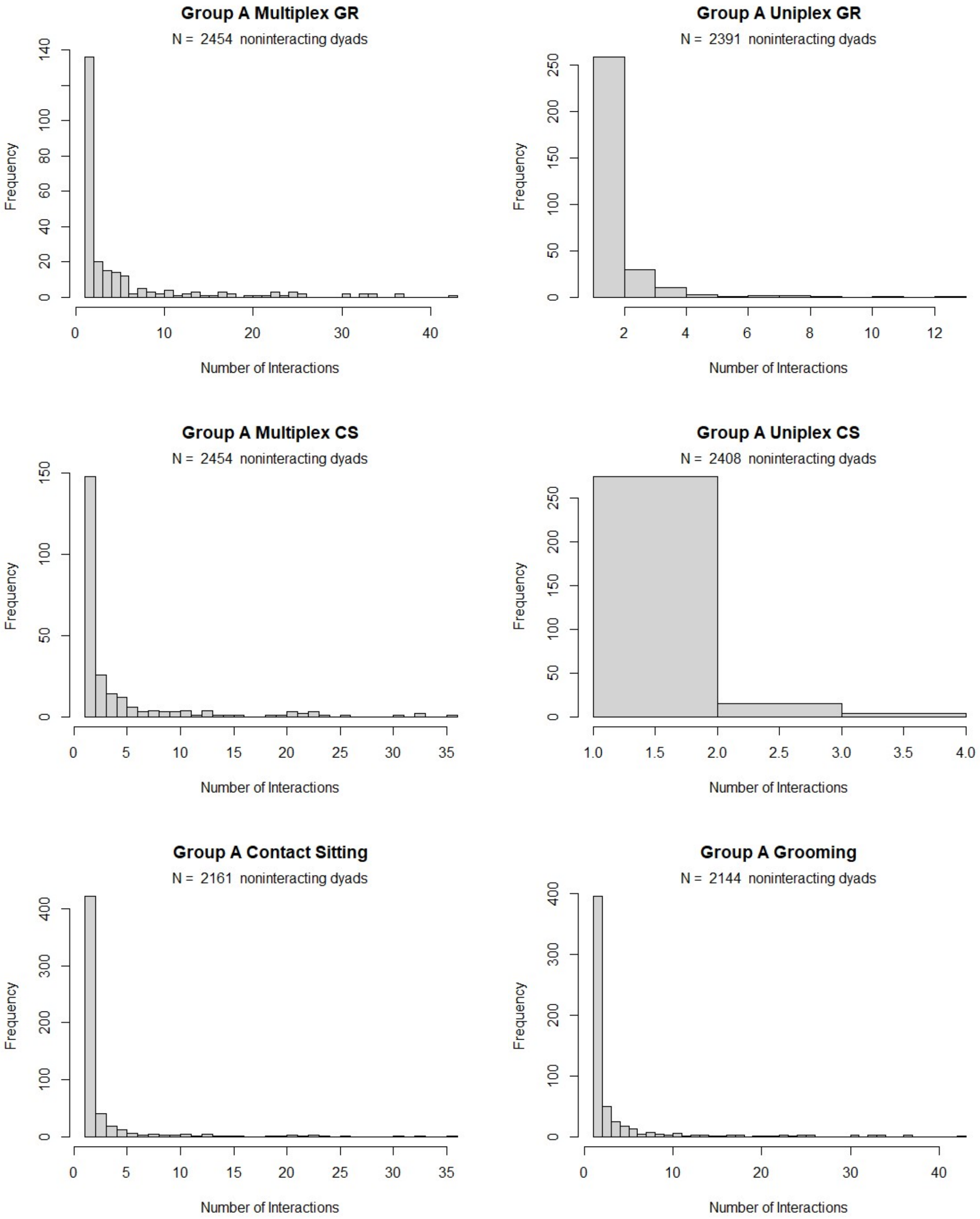

Group B

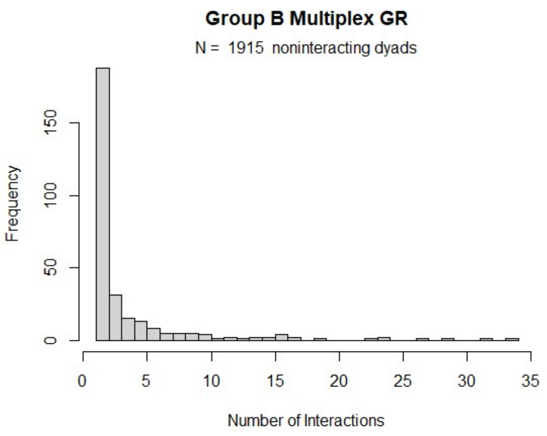
Group B Multiplex GR
N = 1915 noninteracting dyads
Frequency
0
50
100
150
0
5
10
15
20
25
30
35
Number of Interactions

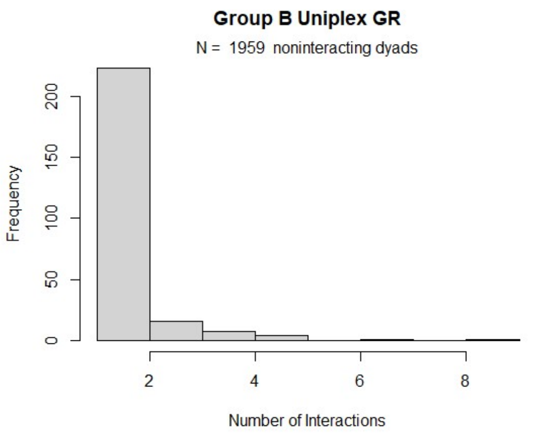
Group B Uniplex GR
N = 1959 noninteracting dyads
Frequency
0
50
100
150
200
2
4
6
8
Number of Interactions

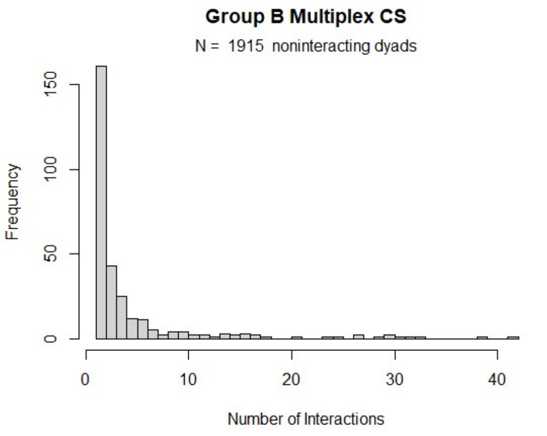
Group B Multiplex CS
N = 1915 noninteracting dyads
Frequency
0
50
100
150
0
10
20
30
40
Number of Interactions

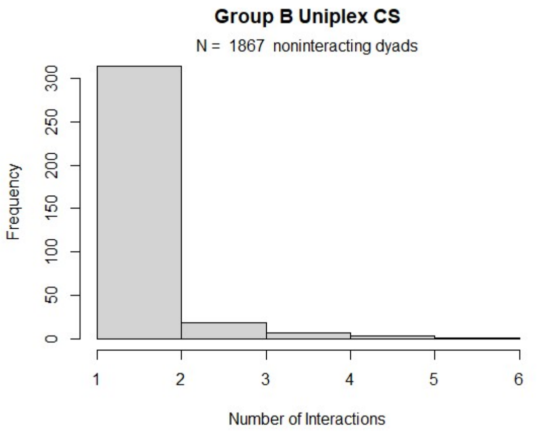
Group B Uniplex CS
N = 1867 noninteracting dyads
Frequency
0
50
100
150
200
250
300
1
2
3
4
5
6
Number of Interactions

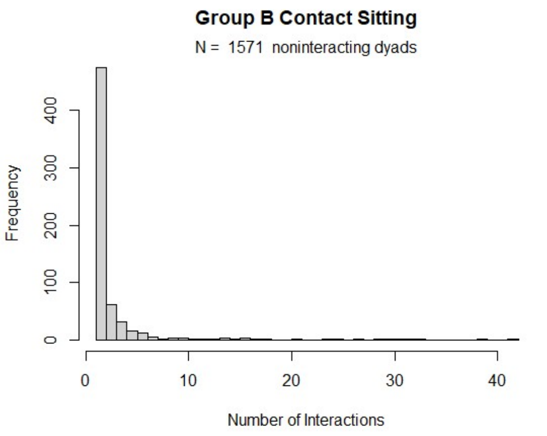
Group B Contact Sitting
N = 1571 noninteracting dyads
Frequency
0
100
200
300
400
0
10
20
30
40
Number of Interactions

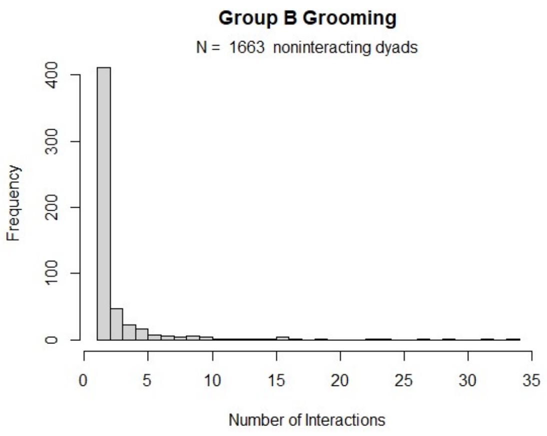
Group B Grooming
N = 1663 noninteracting dyads
Frequency
0
100
200
300
400
0
5
10
15
20
25
30
35
Number of Interactions

Group C

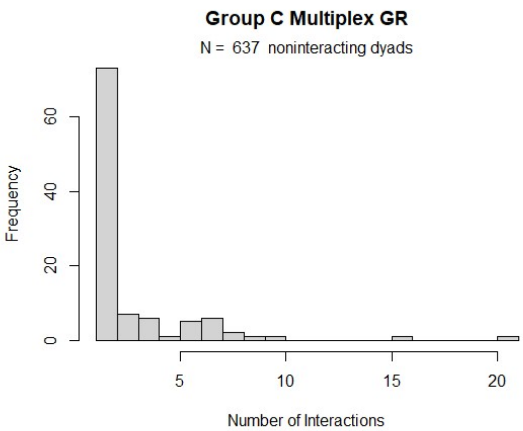
Group C Multiplex GR
N = 637 noninteracting dyads
Frequency
Number of Interactions

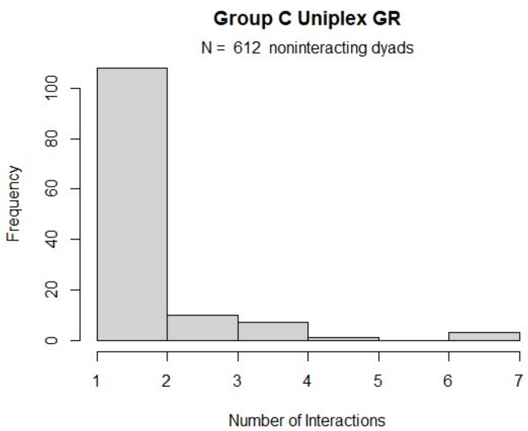
Group C Uniplex GR
N = 612 noninteracting dyads
Frequency
Number of Interactions

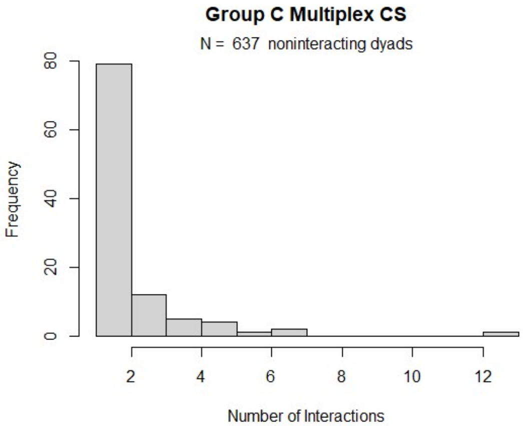
Group C Multiplex CS
N = 637 noninteracting dyads
Frequency
Number of Interactions

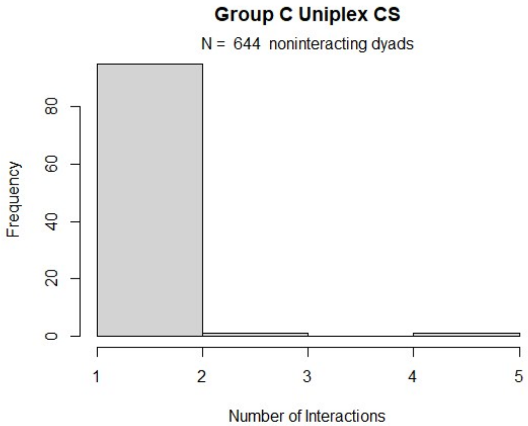
Group C Uniplex CS
N = 644 noninteracting dyads
Frequency
Number of Interactions

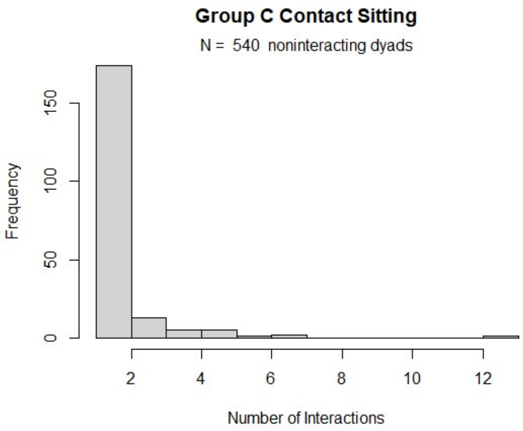
Group C Contact Sitting
N = 540 noninteracting dyads
Frequency
Number of Interactions

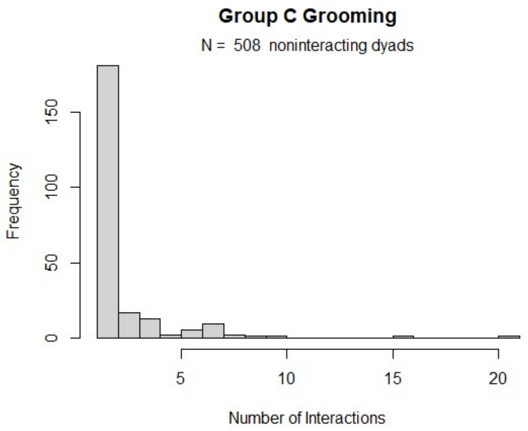
Group C Grooming
N = 508 noninteracting dyads
Frequency
Number of Interactions

Group D

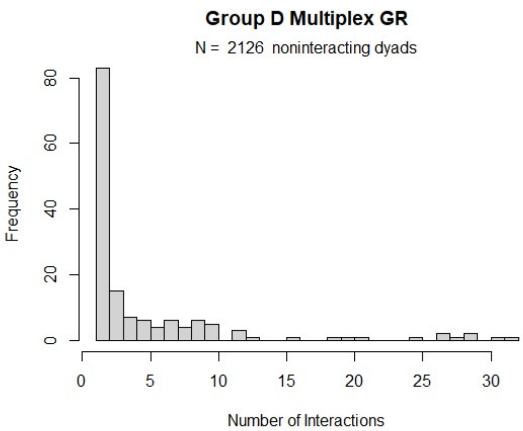
Group D Multiplex GR
N = 2126 noninteracting dyads
Frequency
Number of Interactions

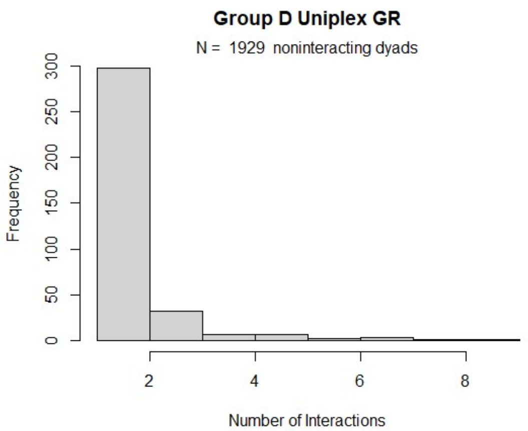
Group D Uniplex GR
N = 1929 noninteracting dyads
Frequency
Number of Interactions

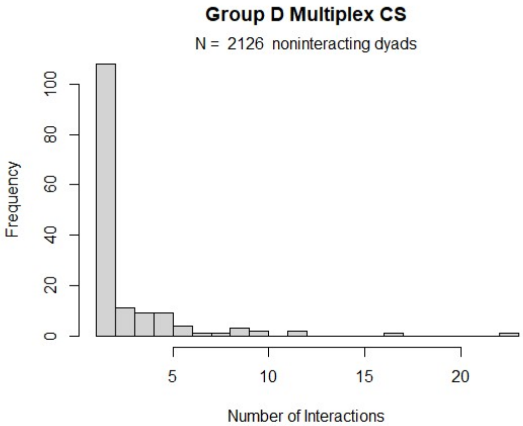
Group D Multiplex CS
N = 2126 noninteracting dyads
Frequency
Number of Interactions

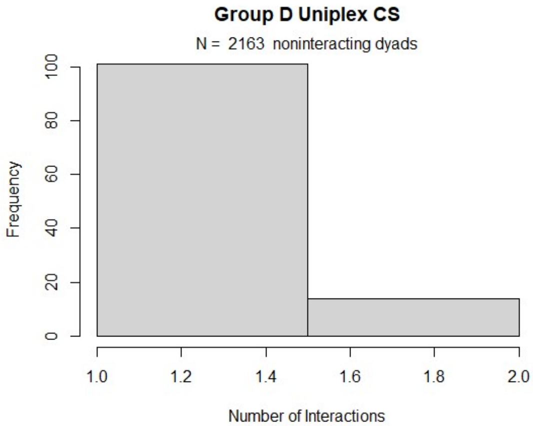
Group D Uniplex CS
N = 2163 noninteracting dyads
Frequency
Number of Interactions

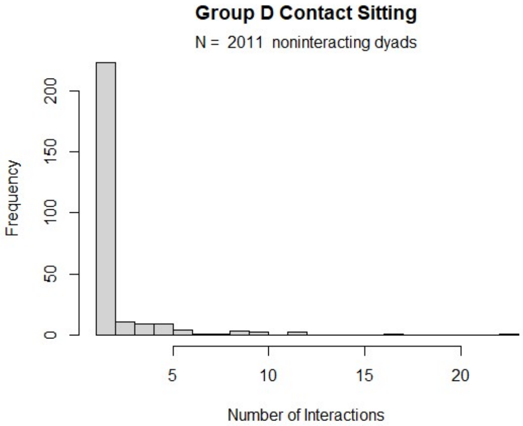
Group D Contact Sitting
N = 2011 noninteracting dyads
Frequency
Number of Interactions

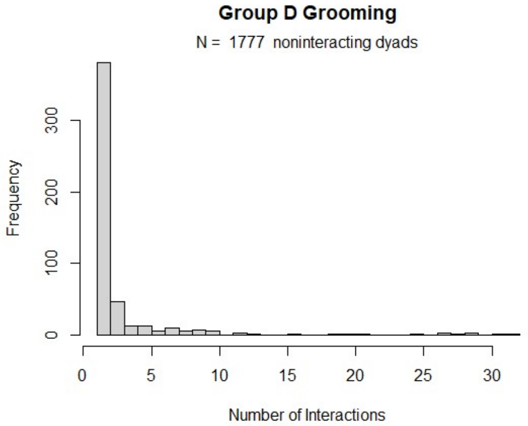
Group D Grooming
N = 1777 noninteracting dyads
Frequency
Number of Interactions

**Figure S2.** Histogram of a) IL-6 and b) TNF-α. Plot a does not include the outlier that was excluded from data analysis.

a) b)

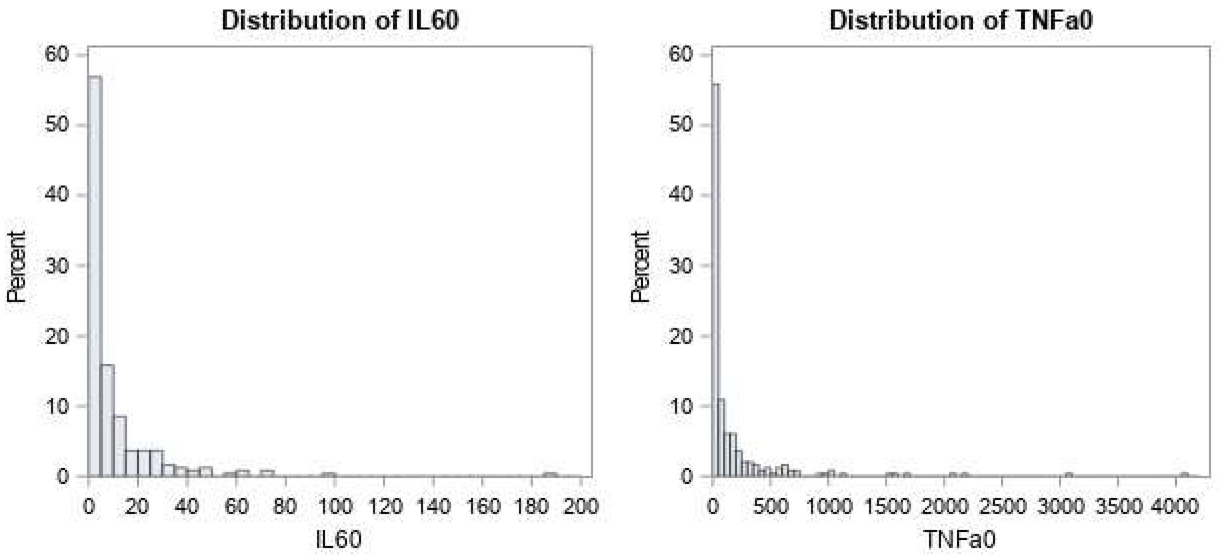


**Figure S3**. Flowchart of the analytical framework

**Step 1: evaluate random effects**

- ID within group

**Step 2: evaluate covariates**

- Rank, dominance certainty, sampling order, age

**Step 3: identify candidate univariate predictors**

- Run univariate statistics on 6 networks with 6 metrics each
  - All groom, Multi groom, Uni groom
  - All contact sit, Multi contact sit, Uni contact sit
  - Select models within lowest AIC of best model
    - If no metrics meet criterion, then test all predictors for those networks in step 4

**Step 4: compare candidate network predictors for paired networks**

- Uni groom vs. Multi groom
- Uni contact sit vs. Multi contact sit
- All groom vs. All contact sit (traditional approach)

**Step 5: report best set of models**

- Report all models with lowest AIC within 2 units of best model

**Supplementary Table 1: Network Node and Edge Details by Group**

| | Multi GR | | Uni GR | | Multi CS | | Uni CS | | All Grooming | | Contact Sit | |
|---|---|---|---|---|---|---|---|---|---|---|---|---|
| **Group** | **N (female)** | **# interaction** | **N (female)** | **# interaction** | **N (female)** | **# interaction** | **N (female)** | **# interaction** | **N (female)** | **# interaction** | **N (female)** | **# interaction** |
| Group A | 74 | 1373 | 73 | 521 | 74 | 1085 | 74 | 372 | 74 | 1894 | 74 | 1457 |
| Group B | 67 | 1094 | 67 | 392 | 67 | 1319 | 67 | 485 | 67 | 1486 | 67 | 1804 |
| Group C | 39 | 288 | 39 | 211 | 39 | 206 | 39 | 114 | 39 | 499 | 39 | 320 |
| Group D | 68 | 776 | 68 | 575 | 68 | 404 | 62 | 129 | 68 | 1351 | 68 | 533 |

**Supplemental Table 3: Model Building Log for IL-6**

| Model | Random | N | # Params | AIC | dAIC |
|---|---|---|---|---|---|
| **1. Establishing the Random Effects Model** | | | | | |
| NULL | Cage | 246 | 1 | 1532.5 | 0 |
| **2. Establishing Relevant Covariates (dAIC is relative to Random Effects Model above)** | | | | | |
| age | Cage | 246 | 2 | 1532.8 | 0.35 |
| dominancecertainty | Cage | 246 | 2 | 1534.4 | 1.97 |
| percentiledominancerank | Cage | 246 | 2 | 1534.5 | 2 |
| percentiledominancerank dominancecertainty percentiledominancerank*dominancecertainty | Cage | 246 | 4 | 1538.31 | 5.81 |
| samplingorder | Cage | 246 | 2 | 1533.5 | 1.04 |
| 3. **Establishing Candidate Predictors (dAIC is relative to Random Effects Model above)** | | | | | |
| MultiGRDegree | Cage | 246 | 2 | 1534.4 | 1.9 |
| MultiGRStrength | Cage | 246 | 2 | 1534.5 | 2.0 |
| MultiGRBetweenness | Cage | 246 | 2 | 1532.9 | 0.5 |
| MultiGREigenvector | Cage | 246 | 2 | 1533.6 | 1.1 |
| MultiGRCloseness | Cage | 246 | 2 | 1534.4 | 1.9 |
| MultiGRClusteringCoefficient | Cage | 246 | 2 | 1532.9 | 0.4 |
| UniGRDegree | Cage | 246 | 2 | 1528.1 | -4.3[b] |
| **UniGRStrength** | **Cage** | **246** | **2** | **1525.3** | **-7.2**[b c] |
| UniGRBetweenness | Cage | 246 | 2 | 1531.3 | -1.2 |
| UniGREigenvector | Cage | 246 | 2 | 1527.8 | -4.7[b] |
| **UniGRCloseness** | **Cage** | **246** | **2** | **1526.0** | **-6.5**[b c] |
| UniGRClusteringCoefficient | Cage | 246 | 2 | 1534.3 | 1.9 |
| MultiCSDegree | Cage | 246 | 2 | 1534.4 | 1.9 |
| MultiCSStrength | Cage | 246 | 2 | 1533.1 | 0.6 |
| MultiCSBetweenness | Cage | 246 | 2 | 1532.9 | 0.5 |
| MultiCSEigenvector | Cage | 246 | 2 | 1533.6 | 1.1 |
| MultiCSCloseness | Cage | 246 | 2 | 1534.4 | 1.9 |
| MultiCSClusteringCoefficient | Cage | 246 | 2 | 1532.9 | 0.4 |
| UniCSDegree | Cage | 240 | 2 | 1492.6 | 1.7 [a] |
| UniCSStrength | Cage | 240 | 2 | 1492.9 | 2.0 [a] |
| UniCSBetweenness | Cage | 240 | 2 | 1491.6 | 0.7 [a] |
| UniCSEigenvector | Cage | 240 | 2 | 1492.7 | 1.8 [a] |
| UniCSCloseness | Cage | 240 | 2 | 1492.5 | 1.6 [a] |
| UniCSClusteringCoefficient | Cage | 240 | 2 | 1491.8 | 0.9 [a] |
| GRDegree | Cage | 246 | 2 | 1530.9 | -1.6 |
| GRStrength | Cage | 246 | 2 | 1532.6 | 0.2 |
| GRBetweenness | Cage | 246 | 2 | 1531.7 | -0.7 |
| GREigenvector | Cage | 246 | 2 | 1529.7 | -2.7[b] |
| GRCloseness | Cage | 246 | 2 | 1531.0 | -1.5 |
| GRClusteringCoefficient | Cage | 246 | 2 | 1533.6 | 1.1 |
| CSDegree | Cage | 246 | 2 | 1534.1 | 1.7 |
| CSStrength | Cage | 246 | 2 | 1533.5 | 1.1 |
| CSBetweenness | Cage | 246 | 2 | 1531.0 | -1.5 |
| CSEigenvector | Cage | 246 | 2 | 1533.5 | 1.0 |
| CSCloseness | Cage | 246 | 2 | 1534.5 | 2.0 |

| | | | | | |
|---|---|---|---|---|---|
| CSClusteringCoefficient | Cage | 246 | 2 | 1532.5 | 0.0 |
| 4. **Final Models (dAIC is relative to Overall Best Model from Step 3 and 4)** | | | | | |
| UniGRDegree MultiGRDegree | Cage | 246 | 3 | 1529.6 | 5.0 |
| UniGRDegree MultiGRStrength | Cage | 246 | 3 | 1530.1 | 5.5 |
| UniGRDegree MultiGRBetweenness | Cage | 246 | 3 | 1529.8 | 5.2 |
| UniGRDegree MultiGREigenvector | Cage | 246 | 3 | 1530.1 | 5.5 |
| UniGRDegree MultiGRCloseness | Cage | 246 | 3 | 1528.3 | 3.7 |
| UniGRDegree MultiGRClustering | Cage | 246 | 3 | 1529.3 | 4.7 |
| **UniGRStrength MultiGRDegree** | **Cage** | **246** | **3** | **1526.5** | **1.9** [c] |
| UniGRStrength MultiGRStrength | Cage | 246 | 3 | 1527.3 | 2.7 |
| UniGRStrength MultiGRBetweenness | Cage | 246 | 3 | 1527.1 | 2.5 |
| UniGRStrength MultiGREigenvector | Cage | 246 | 3 | 1527.3 | 2.7 |
| **UniGRStrength MultiGRCloseness** | **Cage** | **246** | **3** | **1524.9** | **0.3** [c] |
| **UniGRStrength MultiGRClustering** | **Cage** | **246** | **3** | **1526.5** | **1.9** [c] |
| UniGREigenvector MultiGRDegree | Cage | 246 | 3 | 1529.4 | 4.8 |
| UniGREigenvector MultiGRStrength | Cage | 246 | 3 | 1529.8 | 5.2 |
| UniGREigenvector MultiGRBetweenness | Cage | 246 | 3 | 1529.6 | 5.0 |
| UniGREigenvector MultiGREigenvector | Cage | 246 | 3 | 1529.8 | 5.2 |
| UniGREigenvector MultiGRCloseness | Cage | 246 | 3 | 1528.0 | 3.4 |
| UniGREigenvector MultiGRClustering | Cage | 246 | 3 | 1528.7 | 4.1 |
| UniGRCloseness MultiGRDegree | Cage | 246 | 3 | 1527.2 | 2.6 |
| UniGRCloseness MultiGRStrength | Cage | 246 | 3 | 1528.0 | 3.4 |
| UniGRCloseness MultiGRBetweenness | Cage | 246 | 3 | 1528.0 | 3.4 |
| UniGRCloseness MultiGREigenvector | Cage | 246 | 3 | 1528.0 | 3.4 |
| **UniGRCloseness MultiGRCloseness** | **Cage** | **246** | **3** | **1524.6** | **0.0** [c] |
| UniGRCloseness MultiGRClustering | Cage | 246 | 3 | 1527.4 | 2.8 |
| GREigenvector CSDegree | Cage | 246 | 3 | 1531.7 | 7.1 |
| GREigenvector CSStrength | Cage | 246 | 3 | 1530.0 | 5.4 |
| GREigenvector CSBetweenness | Cage | 246 | 3 | 1530.1 | 5.5 |
| GREigenvector CSEigenvector | Cage | 246 | 3 | 1531.7 | 7.1 |
| GREigenvector CSCloseness | Cage | 246 | 3 | 1530.7 | 6.1 |
| GREigenvector CSClustering | Cage | 246 | 3 | 1529.1 | 4.5 |

[a] dAIC compared to random effects only model including only the 240 animals present in the UniCS networks. AIC for this model was 1490.9.

[b] candidate variables for entry into the final models.

[c] **candidate final models**

**Supplemental Table 4: Model Building Log for TNF-α**

| Model | Random | N | # Params | AIC | dAIC |
|---|---|---|---|---|---|
| **1. Establishing the Random Effects Model** | | | | | |
| NULL | Cage | 247 | 1 | 2639.56 | 0 |
| **2. Establishing Covariates (dAIC is relative to Random Effects Model above)** | | | | | |
| age | Cage | 247 | 2 | 2641.32 | 1.76 |
| dominancecertainty | Cage | 247 | 2 | 2640.89 | 1.33 |
| percentiledominancerank | Cage | 247 | 2 | 2641.21 | 1.65 |
| percentiledominancerank dominancecertainty percentiledominancerank*dominancecertainty | Cage | 247 | 4 | 2641.02 | 1.46 |
| samplingorder | Cage | 247 | 2 | 2641.13 | 1.56 |
| **3. Establishing Candidate Predictors (dAIC is relative to Random Effects Model above)** | | | | | |
| MultiGRDegree | Cage | 247 | 2 | 2640.8 | 1.22 |
| MultiGRDegreeWeight | Cage | 247 | 2 | 2641.5 | 1.92 |
| MultiGRBetweenness | Cage | 247 | 2 | 2641.4 | 1.83 |
| MultiGREigenvector | Cage | 247 | 2 | 2641.6 | 1.98 |
| MultiGRCloseness | Cage | 247 | 2 | 2639.7 | 0.14 |
| MultiGRClusteringCoefficient | Cage | 247 | 2 | 2640.2 | 0.63 |
| UniGRDegree | Cage | 247 | 2 | 2636.2 | -3.39 [b] |
| UniGRDegreeWeight | Cage | 247 | 2 | 2635.5 | -4.08 [b] |
| UniGRBetweenness | Cage | 247 | 2 | 2637.1 | -2.45 [b] |
| UniGREigenvector | Cage | 247 | 2 | 2637.2 | -2.43 [b] |
| UniGRCloseness | Cage | 247 | 2 | 2635.9 | -3.68 [b] |
| UniGRClusteringCoefficient | Cage | 247 | 2 | 2641.5 | 1.93 |
| MultiCSDegree | Cage | 247 | 2 | 2640.81 | 1.22 |
| MultiCSStrength | Cage | 247 | 2 | 2640.03 | 0.44 |
| MultiCSBetweenness | Cage | 247 | 2 | 2641.41 | 1.83 |
| MultiCSEigenvector | Cage | 247 | 2 | 2641.56 | 1.98 |
| MultiCSCloseness | Cage | 247 | 2 | 2639.73 | 0.14 |
| MultiCSClusteringCoefficient | Cage | 247 | 2 | 2640.21 | 0.63 |
| UniCSDegree | Cage | 241 | 2 | 2555.90 | -1.60 [a] |
| UniCSStrength | Cage | 241 | 2 | 2554.30 | -3.20 [a b] |
| UniCSBetweenness | Cage | 241 | 2 | 2559.29 | 1.79 [a] |
| UniCSEigenvector | Cage | 241 | 2 | 2556.74 | -0.76 [a] |
| UniCSCloseness | Cage | 241 | 2 | 2552.94 | -4.56 [a b] |
| UniCSClusteringCoefficient | Cage | 241 | 2 | 2559.46 | 1.96 [a] |
| GRDegree | Cage | 247 | 2 | 2640.16 | 0.58 |
| GRStrength | Cage | 247 | 2 | 2640.80 | 1.21 |
| GRBetweenness | Cage | 247 | 2 | 2639.14 | -0.45 |
| GREigenvector | Cage | 247 | 2 | 2639.27 | -0.31 |
| GRCloseness | Cage | 247 | 2 | 2640.19 | 0.60 |
| GRClusteringCoefficient | Cage | 247 | 2 | 2639.86 | 0.28 |
| CSDegree | Cage | 247 | 2 | 2637.92 | -1.67 |
| CSStrength | Cage | 247 | 2 | 2638.44 | -1.14 |
| CSBetweenness | Cage | 247 | 2 | 2641.58 | 2.00 |
| CSEigenvector | Cage | 247 | 2 | 2640.70 | 1.12 |
| CSCloseness | Cage | 247 | 2 | 2637.47 | -2.11 [b] |

| | | | | | |
|---|---|---|---|---|---|
| CSClusteringCoefficient | Cage | 247 | 2 | 2641.41 | 1.83 |
| **4. Final Models (dAIC is relative to the Random Effects Model)** | | | | | |
| UniGRDegree MultiGRDegree | Cage | 247 | 3 | 2633.1 | 3.00 |
| UniGRDegree MultiGRStrength | Cage | 247 | 3 | 2638.0 | 7.90 |
| UniGRDegree MultiGRBetweenness | Cage | 247 | 3 | 2638.2 | 8.10 |
| UniGRDegree MultiGREigenvector | Cage | 247 | 3 | 2636.5 | 6.40 |
| UniGRDegree MultiGRCloseness | Cage | 247 | 3 | 2632.4 | 2.30 |
| UniGRDegree MultiGRClustering | Cage | 247 | 3 | 2634.5 | 4.40 |
| **UniGRStrength MultiGRDegree** | **Cage** | **247** | **3** | **2631.4** | **1.30** [c] |
| UniGRStrength MultiGRStrength | Cage | 247 | 3 | 2637.3 | 7.20 |
| UniGRStrength MultiGRBetweenness | Cage | 247 | 3 | 2637.4 | 7.30 |
| UniGRStrength MultiGREigenvector | Cage | 247 | 3 | 2635.4 | 5.30 |
| **UniGRStrength MultiGRCloseness** | **Cage** | **247** | **3** | **2630.2** | **0.10** [c] |
| UniGRStrength MultiGRClustering | Cage | 247 | 3 | 2632.9 | 2.80 |
| UniGRBetweenness MultiGRDegree | Cage | 247 | 3 | 2634.9 | 4.80 |
| UniGRBetweenness MultiGRStrength | Cage | 247 | 3 | 2639.0 | 8.90 |
| UniGRBetweenness MultiGRBetweenness | Cage | 247 | 3 | 2639.1 | 9.00 |
| UniGRBetweenness MultiGREigenvector | Cage | 247 | 3 | 2637.7 | 7.60 |
| UniGRBetweenness MultiGRCloseness | Cage | 247 | 3 | 2632.3 | 2.20 |
| UniGRBetweenness MultiGRClustering | Cage | 247 | 3 | 2636.1 | 6.00 |
| UniGREigenvector MultiGRDegree | Cage | 247 | 3 | 2635.2 | 5.10 |
| UniGREigenvector MultiGRStrength | Cage | 247 | 3 | 2639.1 | 9.00 |
| UniGREigenvector MultiGRBetweenness | Cage | 247 | 3 | 2639.1 | 9.00 |
| UniGREigenvector MultiGREigenvector | Cage | 247 | 3 | 2637.8 | 7.70 |
| UniGREigenvector MultiGRCloseness | Cage | 247 | 3 | 2633.9 | 3.80 |
| UniGREigenvector MultiGRClustering | Cage | 247 | 3 | 2636.3 | 6.20 |
| **UniGRCloseness MultiGRDegree** | **Cage** | **247** | **3** | **2631.7** | **1.60** [c] |
| UniGRCloseness MultiGRStrength | Cage | 247 | 3 | 2637.8 | 7.70 |
| UniGRCloseness MultiGRBetweenness | Cage | 247 | 3 | 2637.9 | 7.80 |
| UniGRCloseness MultiGREigenvector | Cage | 247 | 3 | 2636.1 | 6.00 |
| **UniGRCloseness MultiGRCloseness** | **Cage** | **247** | **3** | **2630.1** | **0.00** [c] |
| UniGRCloseness MultiGRClustering | Cage | 247 | 3 | 2634.1 | 4.00 |
| GRDegree CSCloseness | Cage | 247 | 3 | 2635.1 | 5.00 |
| GRStrength CSCloseness | Cage | 247 | 3 | 2636.9 | 6.80 |
| GRBetweenness CSCloseness | Cage | 247 | 3 | 2634.1 | 4.00 |
| **GREigenvector CSCloseness** | **Cage** | **247** | **3** | **2631.9** | **1.80** [c] |
| GRCloseness CSCloseness | Cage | 247 | 3 | 2633.3 | 3.20 |
| GRClustering CSCloseness | Cage | 247 | 3 | 2637.5 | 7.40 |
| MultiCSDegree UniCSStrength | Cage | 241 | 3 | 2556.1 | 7.8 [d] |
| MultiCSStrength UniCSStrength | Cage | 241 | 3 | 2556.2 | 7.9 [d] |
| MultiCSBetweenness UniCSStrength | Cage | 241 | 3 | 2555.7 | 7.4 [d] |
| MultiCSEigenvector UniCSStrength | Cage | 241 | 3 | 2556.2 | 7.9 [d] |
| MultiCSCloseness UniCSStrength | Cage | 241 | 3 | 2555.4 | 7.1 [d] |
| MultiCSClustering UniCSStrength | Cage | 241 | 3 | 2555.9 | 7.6 [d] |
| MultiCSDegree UniCSCloseness | Cage | 241 | 3 | 2554.5 | 6.2 [d] |
| MultiCSStrength UniCSCloseness | Cage | 241 | 3 | 2554.8 | 6.5 [d] |
| MultiCSBetweenness UniCSCloseness | Cage | 241 | 3 | 2554.4 | 6.1 [d] |
| MultiCSEigenvector UniCSCloseness | Cage | 241 | 3 | 2554.9 | 6.6 [d] |
| MultiCSCloseness UniCSCloseness | Cage | 241 | 3 | 2554.3 | 6.0 [d] |
| MultiCSClustering UniCSCloseness | Cage | 241 | 3 | 2554.5 | 6.2 [d] |

[a] dAIC compared to random effects only model including only the 241 animals present in the UniCS networks. AIC for this model was 2557.5.

[b] candidate variables for entry into the final models.

**[c] candidate final models**

[d] dAIC compared to the overall best fit model [c] including MultiGRcloseness and UniGRcloseness for only the 241 animals present in the UniCS network in step 4. AIC for this reduced model was 2548.3.
