## SupplementaryTable2 for "Differential effects of multiplex and uniplex affiliative relationships on biomarkers of inflammation"

Supplementary Table S2

| | |
|---|---|
| CS | Contact Sitting |
| GR | Grooming |
| Multi | Multiplex |
| Uni | Uniplex |

| n = 247 | GRdegree | GRstrength | GRbetweenness | GReigenvector | GRcloseness | GRclustering | CSdegree | CSstrength | CSbetweenness | CSeigenvector | CScloseness | CSclustering |
|---|---|---|---|---|---|---|---|---|---|---|---|---|
| **GRdegree** | 1 | 0.54196 | 0.76404 | 0.85181 | 0.80626 | -0.36529 | 0.45231 | 0.27427 | 0.23513 | 0.34793 | 0.28143 | 0.00792 |
| **GRstrength** | 0.54196 | 1 | 0.35184 | 0.35529 | 0.30197 | -0.2194 | 0.28415 | 0.38112 | 0.03714 | 0.07087 | 0.11073 | -0.02248 |
| **GRbetweenness** | 0.76404 | 0.35184 | 1 | 0.79445 | 0.83473 | -0.38849 | 0.198 | 0.03402 | 0.29556 | 0.28566 | 0.22284 | -0.02359 |
| **GReigenvector** | 0.85181 | 0.35529 | 0.79445 | 1 | 0.93622 | -0.13998 | 0.26688 | 0.02536 | 0.35579 | 0.43797 | 0.28525 | 0.0501 |
| **GRcloseness** | 0.80626 | 0.30197 | 0.83473 | 0.93622 | 1 | -0.23396 | 0.3361 | 0.09136 | 0.35942 | 0.48771 | 0.41232 | 0.08185 |
| **GRclustering** | -0.36529 | -0.2194 | -0.38849 | -0.13998 | -0.23396 | 1 | -0.08976 | -0.05951 | -0.11818 | 0.02707 | 0.05802 | 0.25034 |
| **CSdegree** | 0.45231 | 0.28415 | 0.198 | 0.26688 | 0.3361 | -0.08976 | 1 | 0.79332 | 0.32131 | 0.77249 | 0.84918 | 0.10224 |
| **CSstrength** | 0.27427 | 0.38112 | 0.03402 | 0.02536 | 0.09136 | -0.05951 | 0.79332 | 1 | 0.04524 | 0.4779 | 0.64683 | 0.16318 |
| **CSbetweenness** | 0.23513 | 0.03714 | 0.29556 | 0.35579 | 0.35942 | -0.11818 | 0.32131 | 0.04524 | 1 | 0.58202 | 0.30277 | -0.42841 |
| **CSeigenvector** | 0.34793 | 0.07087 | 0.28566 | 0.43797 | 0.48771 | 0.02707 | 0.77249 | 0.4779 | 0.58202 | 1 | 0.78616 | 0.09438 |
| **CScloseness** | 0.28143 | 0.11073 | 0.22284 | 0.28525 | 0.41232 | 0.05802 | 0.84918 | 0.64683 | 0.30277 | 0.78616 | 1 | 0.19415 |
| **CSclustering** | 0.00792 | -0.02248 | -0.02359 | 0.0501 | 0.08185 | 0.25034 | 0.10224 | 0.16318 | -0.42841 | 0.09438 | 0.19415 | 1 |

| n = 247 | MultiGRdegree | MultiGRstrength | MultiGRbetweenness | MultiGReigenvector | MultiGRcloseness | MultiGRclustering | UniGRdegree | UniGRstrength | UniGRbetweenness | UniGReigenvector | UniGRcloseness | UniGRclustering |
|---|---|---|---|---|---|---|---|---|---|---|---|---|
| **MultiGRdegree** | 1 | 0.43905 | 0.46609 | 0.80655 | 0.81845 | -0.2011 | 0.18575 | 0.1885 | 0.2284 | 0.21137 | 0.08414 | -0.09546 |
| **MultiGRstrength** | 0.43905 | 1 | 0.12229 | 0.35894 | 0.23732 | 0.08934 | 0.04257 | 0.07467 | -0.00525 | -0.05811 | -0.10023 | -0.12182 |
| **MultiGRbetweenness** | 0.46609 | 0.12229 | 1 | 0.3966 | 0.33487 | -0.33554 | 0.28305 | 0.27929 | 0.20786 | 0.28622 | 0.35913 | 0.02743 |
| **MultiGReigenvector** | 0.80655 | 0.35894 | 0.3966 | 1 | 0.68815 | -0.04707 | 0.12761 | 0.16872 | 0.1689 | 0.13494 | 0.09928 | 0.00553 |
| **MultiGRcloseness** | 0.81845 | 0.23732 | 0.33487 | 0.68815 | 1 | -0.22832 | -0.02858 | -0.00692 | 0.23869 | 0.10753 | -0.03906 | -0.04866 |
| **MultiGRclustering** | -0.2011 | 0.08934 | -0.33554 | -0.04707 | -0.22832 | 1 | -0.15183 | -0.13227 | -0.16603 | -0.12301 | -0.14177 | 0.11837 |
| **UniGRdegree** | 0.18575 | 0.04257 | 0.28305 | 0.12761 | -0.02858 | -0.15183 | 1 | 0.87333 | 0.69319 | 0.9026 | 0.87904 | 0.06162 |
| **UniGRstrength** | 0.1885 | 0.07467 | 0.27929 | 0.16872 | -0.00692 | -0.13227 | 0.87333 | 1 | 0.61678 | 0.78856 | 0.77245 | 0.06929 |
| **UniGRbetweenness** | 0.2284 | -0.00525 | 0.20786 | 0.1689 | 0.23869 | -0.16603 | 0.69319 | 0.61678 | 1 | 0.68407 | 0.70208 | -0.0983 |
| **UniGReigenvector** | 0.21137 | -0.05811 | 0.28622 | 0.13494 | 0.10753 | -0.12301 | 0.9026 | 0.78856 | 0.68407 | 1 | 0.89482 | 0.1947 |
| **UniGRcloseness** | 0.08414 | -0.10023 | 0.35913 | 0.09928 | -0.03906 | -0.14177 | 0.87904 | 0.77245 | 0.70208 | 0.89482 | 1 | 0.14048 |
| **UniGRclustering** | -0.09546 | -0.12182 | 0.02743 | 0.00553 | -0.04866 | 0.11837 | 0.06162 | 0.06929 | -0.0983 | 0.1947 | 0.14048 | 1 |

| n (Multi CS) = 247<br>n (UniCS) = 241 | MultiCSdegree | MultiCSstrength | MultiCSbetweenness | MultiCSeigenvector | MultiCScloseness | MultiCSclustering | UniCSdegree | UniCSstrength | UniCSbetweenness | UniCSeigenvector | UniCScloseness | UniCSclustering |
|---|---|---|---|---|---|---|---|---|---|---|---|---|
| **MultiCSdegree** | 1 | 0.58019 | 0.46609 | 0.80655 | 0.81845 | -0.2011 | 0.39007 | 0.42482 | -0.08454 | 0.23034 | 0.43507 | 0.18807 |
| **MultiCSstrength** | 0.58019 | 1 | 0.01509 | 0.39156 | 0.52144 | -0.04738 | 0.51612 | 0.52101 | -0.13585 | 0.25165 | 0.52271 | 0.27831 |
| **MultiCSbetweenness** | 0.46609 | 0.01509 | 1 | 0.3966 | 0.33487 | -0.33554 | -0.05667 | -0.06007 | 0.20426 | 0.02139 | -0.08596 | -0.1425 |
| **MultiCSeigenvector** | 0.80655 | 0.39156 | 0.3966 | 1 | 0.68815 | -0.04707 | 0.24684 | 0.25705 | -0.04517 | 0.19825 | 0.32141 | 0.08602 |
| **MultiCScloseness** | 0.81845 | 0.52144 | 0.33487 | 0.68815 | 1 | -0.22832 | 0.4949 | 0.51407 | -0.04828 | 0.40659 | 0.68 | 0.32777 |
| **MultiCSclustering** | -0.2011 | -0.04738 | -0.33554 | -0.04707 | -0.22832 | 1 | -0.16256 | -0.14298 | -0.15128 | -0.15848 | -0.17952 | -0.05588 |
| **UniCSdegree** | 0.39007 | 0.51612 | -0.05667 | 0.24684 | 0.4949 | -0.16256 | 1 | 0.96314 | 0.3578 | 0.8297 | 0.86695 | 0.25874 |
| **UniCSstrength** | 0.42482 | 0.52101 | -0.06007 | 0.25705 | 0.51407 | -0.14298 | 0.96314 | 1 | 0.2981 | 0.77105 | 0.83503 | 0.24948 |
| **UniCSbetweenness** | -0.08454 | -0.13585 | 0.20426 | -0.04517 | -0.04828 | -0.15128 | 0.3578 | 0.2981 | 1 | 0.5844 | 0.22707 | -0.1615 |
| **UniCSeigenvector** | 0.23034 | 0.25165 | 0.02139 | 0.19825 | 0.40659 | -0.15848 | 0.8297 | 0.77105 | 0.5844 | 1 | 0.7879 | 0.26459 |
| **UniCScloseness** | 0.43507 | 0.52271 | -0.08596 | 0.32141 | 0.68 | -0.17952 | 0.86695 | 0.83503 | 0.22707 | 0.7879 | 1 | 0.42021 |
| **UniCSclustering** | 0.18807 | 0.27831 | -0.1425 | 0.08602 | 0.32777 | -0.05588 | 0.25874 | 0.24948 | -0.1615 | 0.26459 | 0.42021 | 1 |
